# Accumulation of TERT in Mitochondria Shows Two Opposing Effects on Apoptosis

**DOI:** 10.1101/2021.12.21.473585

**Authors:** Hiroshi Ebata, Tomohiro Shima, Ryo Iizuka, Sotaro Uemura

**Author notes:** Corresponding authors T.S., S.U.

## Abstract

Telomerase reverse transcriptase (TERT) is a protein that catalyzes the reverse transcription of telomere elongation. TERT is also expected to play a noncanonical role beyond telomere lengthening since it localizes not only in the nucleus but also in mitochondria, where telomeres do not exist. Several studies have reported that mitochondrial TERT regulates apoptosis induced by oxidative stress. However, there remains controversy about whether mitochondrial TERT promotes or inhibits apoptosis, mainly due to the lack of information on changes in the TERT distribution in individual cells over time. Here we simultaneously detected apoptosis and TERT localization after oxidative stress in individual HeLa cells by live-cell tracking. This tracking revealed that the stress-induced accumulation of TERT in mitochondria caused apoptosis but that the accumulation positively correlated with the time until cell death. The results suggest a new model in which mitochondrial TERT has two opposing effects at different stages of apoptosis: it predetermines apoptosis at the first stage of cell-fate determination but also delays apoptosis at the second stage. Because these distinct effects respectively support both sides of the controversy regarding the role of mitochondrial TERT in apoptosis, our model integrates two opposing hypotheses. Furthermore, detailed statistical analysis of TERT mutations, which have been predicted to inhibit TERT transport to mitochondria, revealed that these mutations suppress apoptosis independent of the mitochondrial localization of TERT. Together, these results indicate that the non-canonical functions of TERT affect a wide range of pathways.

## INTRODUCTION

Telomerase reverse transcriptase (TERT) is a protein subunit of the telomerase complex, which elongates the telomeric repeat sequences at chromosomal ends and prevents telomere loss [1]. In humans, most cancer cells express high levels of TERT, whereas somatic cells suppress TERT expression [2]. Since TERT increases the number of cell divisions, it is believed to promote the unlimited growth of cancer cells. Interestingly, several studies have shown that TERT is localized not only in the nucleus but also in mitochondria, which lack telomeric regions [3–5]. Oxidative stress, which eventually leads to cell apoptosis, has been reported to increase TERT localization in mitochondria [4, 5], raising the possibility of a noncanonical role of mitochondrial TERT in apoptosis beyond its telomere elongation function. Mutagenesis studies have proposed that mitochondrial TERT induces apoptosis [4, 6]. However, TERT overexpression has been reported to increase TERT in mitochondria and cell survival after oxidative stress, suggesting that mitochondrial TERT suppresses apoptosis [5, 7]. These reports, while conflicting, have shed light on the possibility that mitochondrial TERT regulates apoptosis.

These conflicting observations can be attributed mainly to three reasons. First, the relationship between TERT localization and cell death in individual cells has not been fully tested. Due to fluorescently labeled TERT failing to retain normal TERT enzymatic activity and cellular distribution, previous studies have only detected mitochondrial TERT after cell fixation or cell disruption [4–6]. Therefore, the eventual fate of each individual cell is unknown. Secondly, there is little temporal information. Previous immunofluorescence- and flow cytometry-based measurements only provide information on cell death and TERT localization at specific times. However, oxidative stress induces cell death through several different pathways [8, 9]. Even within the same pathway, the response time to cell death-inducing stimuli varies among cells. Thus, directly testing the relationship of mitochondrial TERT and cell death requires tracking the two factors over time. Thirdly, classical experimental methods can damage the cells. Immunofluorescence, for example, requires multiple washes for each step of fixation, permeabilization, and antibody binding. Cells undergoing cell death lose their adhesiveness to the dish surface, resulting in some being lost by the multiple washes. The loss of dead cells results in underestimating apoptosis. In flow cytometry measurements, flowing cells are exposed to physical stress, including high pressures, shear forces, electrical charges, shock forces, and rapid temperature changes. These additional stresses can induce stress responses by the cells, resulting in more cell death. Therefore, directly testing the relationship between mitochondrial localization and cell death using stress-free methods is needed.

In response, here we combined imaging-based dead-cell detection methods with live-cell fluorescence imaging of TERT to directly assess the relationship between the mitochondrial localization of TERT and apoptosis of individual cells. Live-cell imaging provides spatial and temporal information of the TERT distribution until cell death. This approach has several advantages over conventional dead-cell detection methods. Namely, live-cell imaging avoids extra physical stress to the cells because it requires only one medium exchange to cause oxidative stress without any washing steps. Also, to visualize TERT distribution in a cell, we fused TERT to a fluorescent protein, mVenus, at a position that did not interfere with the mitochondrial localization or enzymatic activity. This setup enabled us to track the temporal changes of TERT localization. Consequently, we found that the accumulation of TERT in mitochondria emerged immediately after oxidative stress in a subset of cells to cause apoptosis but, at the same time, the accumulation positively correlated with a longer time until cell death. These results suggest that mitochondrial TERT plays distinct roles at different stages of apoptosis. We also elucidated the effects of previously reported mutations in TERT and found that they are independent of TERT localization in mitochondria.

## RESULTS

### TERT mutations R3E/R6E and Y707F do not change TERT mitochondrial localization in HeLa cells

We first engineered TERT mutants and assessed theirs and wild-type TERT localization to mitochondria. We prepared stable HeLa cell lines expressing wild-type TERT and TERT mutants using the Sleeping Beauty system [10]. The R3E/R6E mutation has been reported to inhibit the mitochondrial translocation of TERT due to the location of these residues in the mitochondrial targeting signal (MTS) of TERT [4]. Y707F has been reported to inhibit the oxidative stress-stimulated nuclear export of TERT because phosphorylation of the tyrosine residue by Src kinases correlates with the export [6]. We examined the localization of these mutants by immunofluorescence (Fig. 1A). The degree of TERT mitochondrial localization was quantified by Manders’ colocalization coefficient (MCC) from the immunofluorescence images of these cell lines cultured under normal conditions (no oxidative stress). MCC is an intuitive and direct metric measuring the co-occurrence of the quantity of interest [11]. MCC of R3E/R6E and Y707F TERT mutants and mitochondria approximated that of wild-type TERT (Fig. 1B). This result suggests that these mutations did not prevent TERT transport to mitochondria in normal conditions. Previous studies also evaluated the localization of these TERT mutants but by representative images instead of statistical tests [4, 6]. The difference in localization analysis might be the reason for the different localization patterns observed in ours and previous studies.

**Figure 1.**
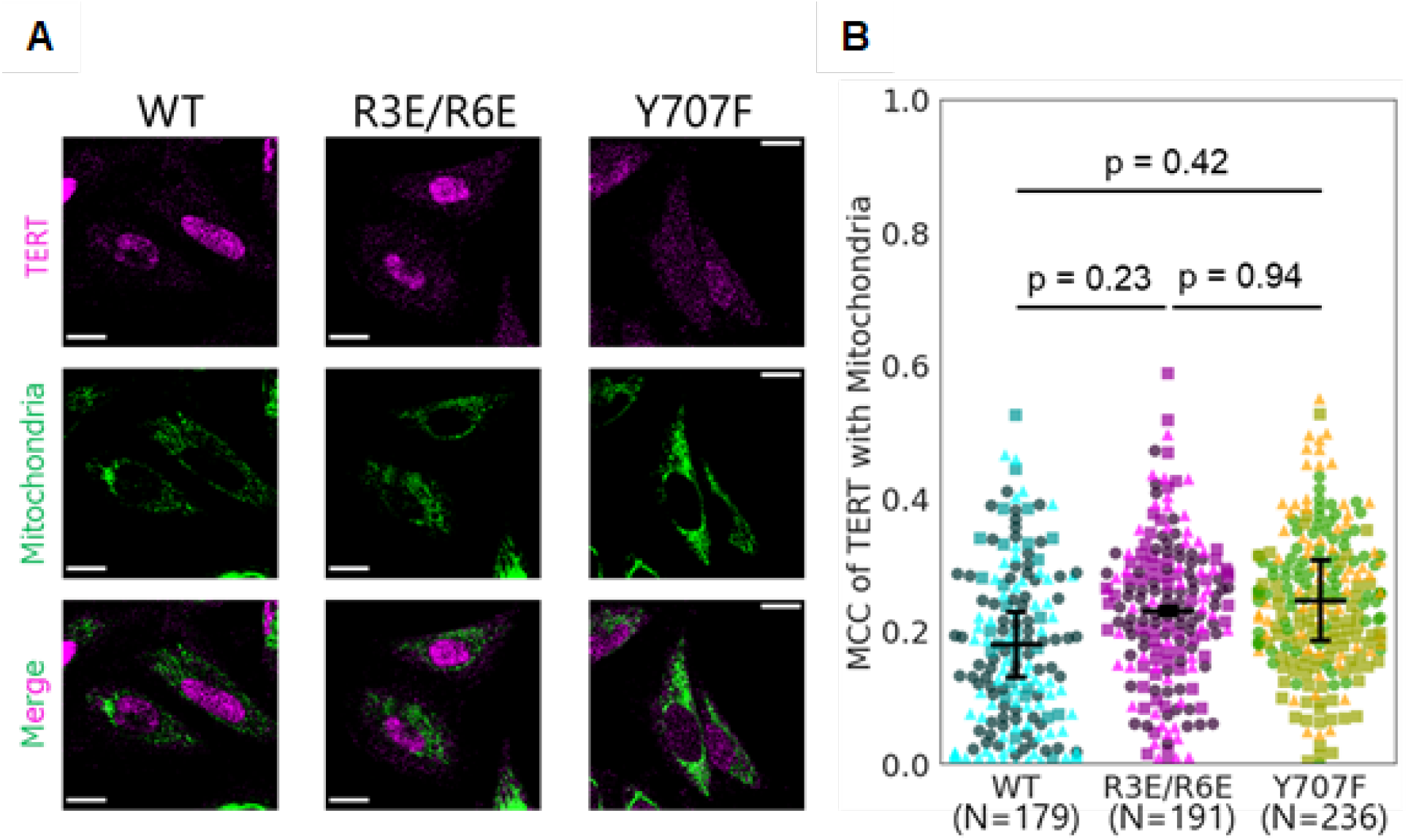
TERT mutations R3E/R6E and Y707F do not change mitochondrial TERT localization in HeLa cells. A. Representative immunofluorescence images of HeLa cells expressing the TERT constructs. Magenta, anti-TERT immunofluorescence; green, MitoTracker Deep Red FM fluorescence. Scale bars, 20 μm. B. Quantification of the mitochondrial localization of TERT. Manders’ colocalization coefficient (MCC) of TERT with mitochondria was calculated from the fluorescence intensity of TERT and mitochondria. Dots show MCC of each cell, and bars show the mean ± 95% C.I. (1.96 SEM) from 3 independent experiments. Different markers represent different experiments. Wild-type (WT): N=179 cells; R3E/R6E: N=191 cells; Y707F: N=236 cells. Steel-Dwass test was performed.

### TERT mutations R3E/R6E and Y707F increase cell survival

Next, to observe the effect of the TERT mutations on the apoptotic response, we introduced oxidative stress to the cells and tracked each cell by live-cell imaging under a fluorescence microscope (Fig. 2A). Conventionally, oxidative stress treatment omits FBS in the medium since FBS scavenges hydrogen peroxide and attenuates the effect of oxidative stress. However, even when we cultured cells in the medium without oxidative stress and FBS, the majority of cells were detached from the dish surface after medium exchange. In addition, even with FBS, oxidative stress treatment caused approximately 70% of the cells expressing wild-type TERT to undergo cell death. Thus, we decided to put oxidative stress to cells with FBS in this work (see Methods Dead-cell detection with live-cell imaging). We determined the time until cell death by the fluorescence intensity of SYTOX Orange, which stains dead cells, and YO-PRO-1, which stains apoptotic cells (Fig. 2B, see Methods Microscopy for live-cell imaging and tracking). Survival curves of these cell lines demonstrated that HeLa cells expressing either TERT mutant were more resistant to oxidative stress than those expressing wild-type TERT (Fig. 2C). This result agrees with previous studies that reported the mutants increased cell viability after oxidative stress [4, 6]. In all conditions, the cell survival percentage reached a plateau at approximately 40 hours after oxidative stress treatment. Finally, the survival curve of the SYTOX Orange signal lagged that of the YO-PRO-1 signal by several hours (Supplementary Fig. S1A).

**Figure 2.**
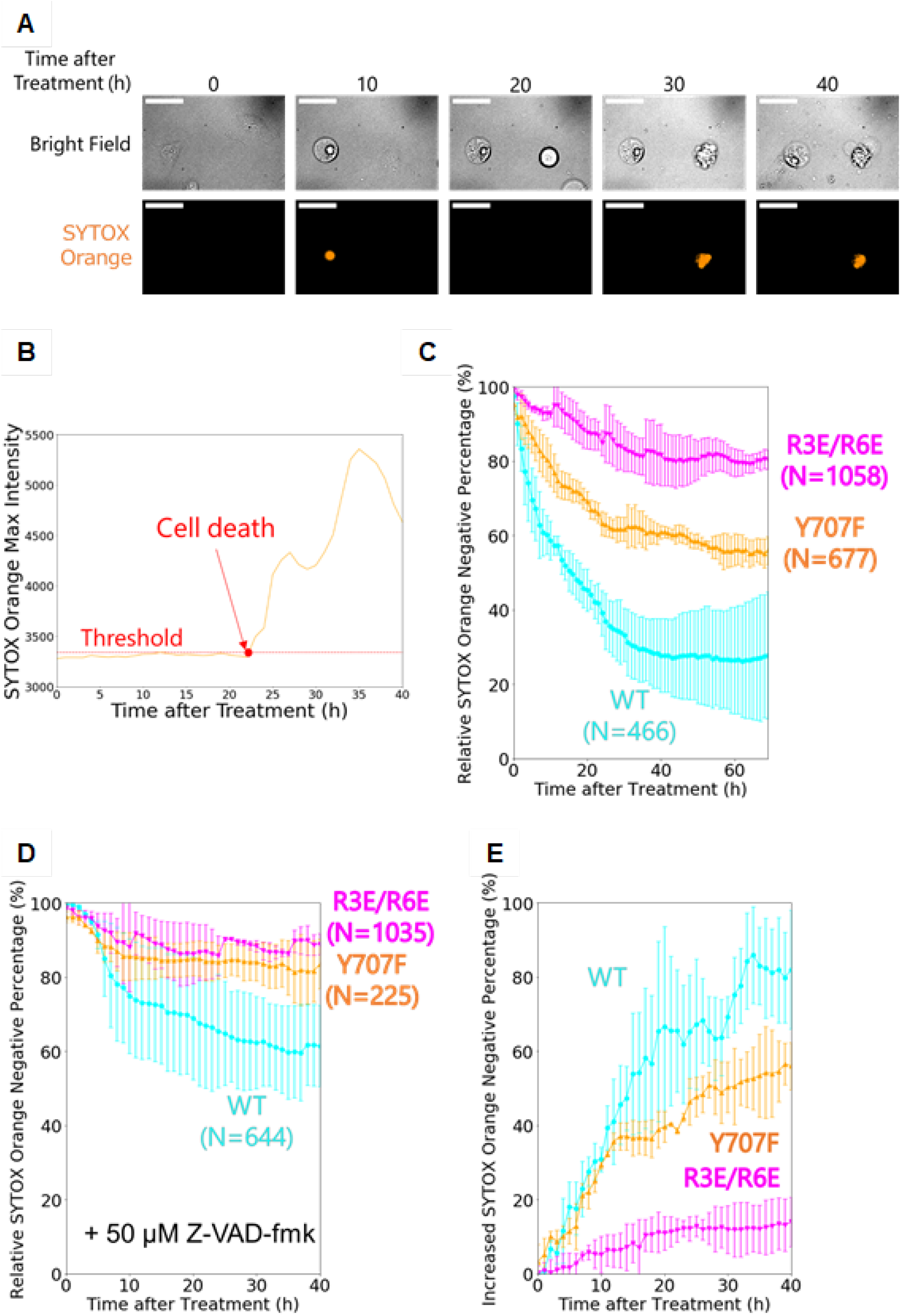
TERT mutations R3E/R6E and Y707F increased the cell survival. A. Representative live-cell images of HeLa cells expressing the TERT constructs. Cells were treated with 267 μM calcium percarbonate for 3 hours before imaging. Orange, SYTOX Orange fluorescence. Scale bars, 50 μm. B. A representative time-course trace of the fluorescence intensity of the dead-cell staining dye SYTOX Orange from the right cell in A. Cells were determined as positive after the SYTOX Orange fluorescence intensity exceeded the threshold (red dashed line). C. Percentage of SYTOX Orange-negative cells after oxidative stress. All percentages were corrected by control experiments without oxidative stress (Supplementary Fig. 1B). Error bars show 95% C.I. (1.96 SEM) from 3 independent experiments. Wild-type (WT): N=466 cells; R3E/R6E: N=1058 cells; Y707F: N=677 cells. D. Percentage of SYTOX Orange-negative cells after oxidative stress when 50 μM Z-VAD-fmk was added. All percentages were corrected by control experiments without oxidative stress (Supplementary Fig. 1B). Error bars show 95% C.I. (1.96 SEM) from 3 independent experiments. Wild-type (WT): N=644 cells; R3E/R6E: N=1035 cells; Y707F: N=225 cells. E. The percentage of SYTOX Orange-negative cells increased by Z-VAD-fmk after oxidative stress. Error bars show 95% C.I. (1.96 SEM) from 3 independent experiments.

To determine whether apoptosis inhibition changes the survival curves, we did the same experiments using a pan-caspase inhibitor, Z-VAD-fmk [12]. Z-VAD-fmk increased the survival of all HeLa cell lines (Fig. 2C), but the extent of the increase differed among the lines (Fig. 2D). Drastic changes in the viability of cells expressing wild-type TERT by Z-VAD-fmk indicated that cell death induced by oxidative stress was predominantly apoptosis, as previously reported [13]. Z-VAD-fmk increased the viability at 40 hours after oxidative stress treatment by approximately 80% in HeLa cells expressing wild-type TERT, 60% in cells expressing TERT with Y707F, and only 20% in cells expressing TERT with R3E/R6E. These results indicate that the caspase inhibition effect of Z-VAD-fmk was blunt in cell lines expressing the TERT mutants, supporting the notion that these mutations inhibit the apoptotic process even without Z-VAD-fmk. Collectively, the lag of the SYTOX Orange signal compared with the YO-PRO-1 signal and the increase in cell survival by Z-VAD-fmk suggest that apoptosis is the predominant cell-death pathway of HeLa cells under oxidative stress and that the mutations inhibit apoptosis.

### Probing TERT without interfering with its activity or mitochondrial localization

Next, to directly investigate the relationship between the mitochondrial localization of TERT and apoptosis in individual cells, we inserted mVenus into TERT. In preparation for the visualization of TERT localization by live-cell tracking, we assessed the effect of the insertion on the telomerase activity and localization pattern of TERT. Conventionally, epitope tags or fluorescent proteins are conjugated to TERT at the N- or C-terminus [6, 14, 15]. However, TERT has an MTS at the N-terminus and an essential domain for its telomerase enzymatic activity at the C-terminus [3, 16, 17], suggesting conjugation at either location could affect TERT function. Moreover, a previous report suggested that mitochondrial TERT functions dependent on its reverse transcriptase activity associated with mitochondrial DNA and tRNAs [18], which is consistent with other reports that TERT can bind to mitochondrial nucleotides [19, 20]. Therefore, we inserted mVenus between A67 and A68 (hereafter mVenus-TERT), which we considered an ideal location because it is far from the reaction site of the telomerase complex and telomerase co-factors (Fig. 3A) [21, 22]. We evaluated the telomerase activity and localization of mVenus-TERT by qPCR-based telomerase activity assay and immunofluorescence (Fig. 3B-D). As expected, mVenus insertion did not interfere with the telomerase activity, while conjugating mVenus to the C-terminus (TERT-mVenus) deteriorated telomerase activity (Fig. 3B). mVenus-TERT showed a higher mitochondrial localization, which might result from inhibiting nuclear import of TERT, but at least, mVenus insertion did not decrease mitochondrial localization of TERT. Accordingly, we used mVenus-TERT to visualize TERT during apoptosis by live-cell imaging.

**Figure 3.**
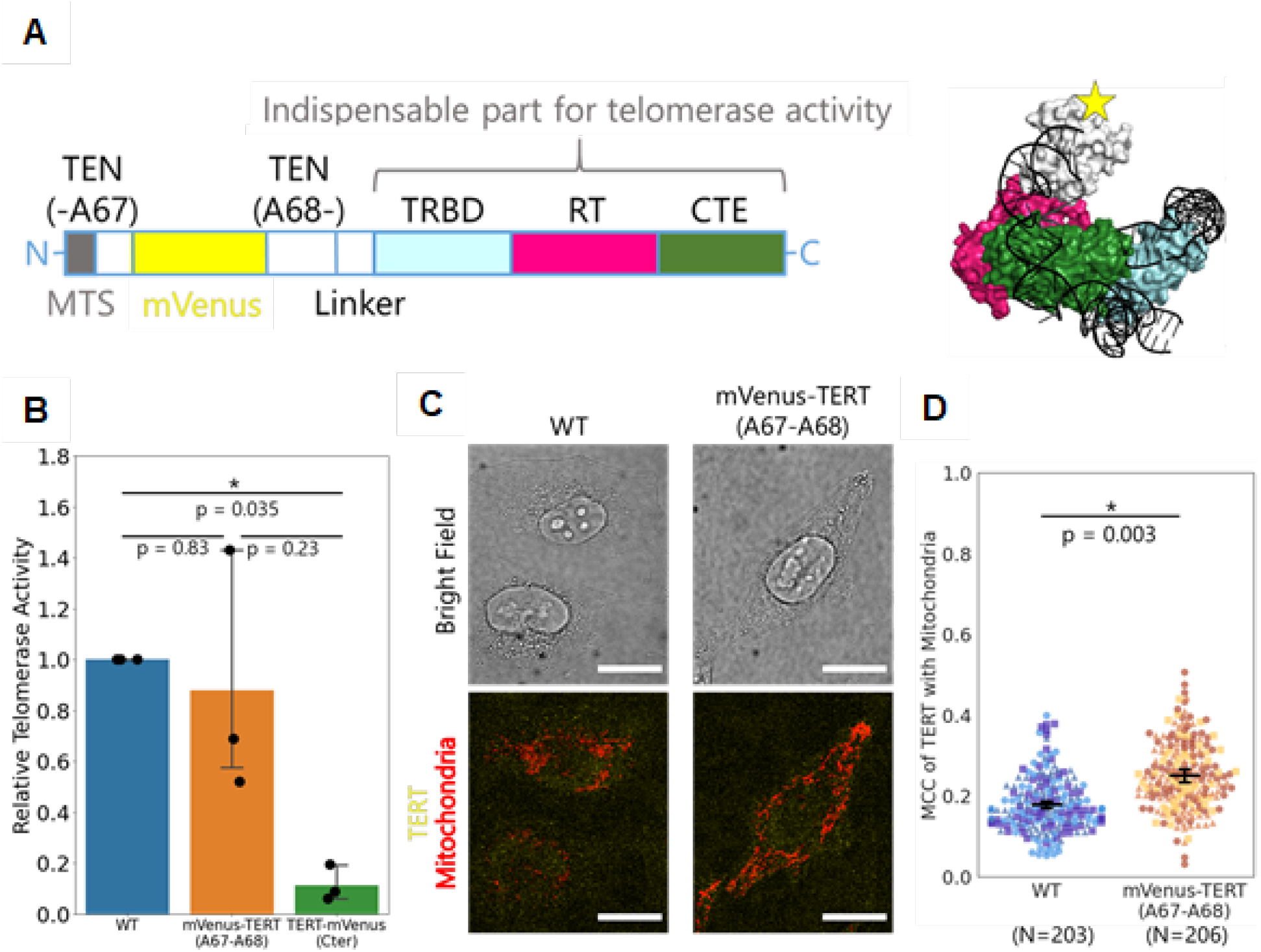
TERT was probed without interfering its activity or mitochondrial localization. A. Schematic representation of mVenus-TERT for live-cell tracking. The structural model was created from EMD-7518. MTS, mitochondrial targeting signal; TEN, telomerase essential N-terminal domain; TRBD, telomerase RNA binding domain; RT, reverse transcriptase domain; CTE, C-terminal extension. B. Telomerase activity of whole cell extracts from HeLa cells expressing the TERT constructs. TERT activity was measured by a qPCR-based TRAP assay. Dots show each data, and bars show the mean ± 95% C.I. (1.96 SEM) from 3 independent experiments. Steel-Dwass test was performed. C. Representative immunofluorescence images of HeLa cells expressing the TERT constructs. Yellow, anti-TERT immunofluorescence; red, MitoTracker Deep Red FM fluorescence. Scale bars, 20 μm. D. Quantification of the TERT localization. MCC of TERT with mitochondria was calculated from the TERT and mitochondria fluorescence intensities. Dots show MCC of each cell, and bars show the mean ± 95% C.I. (1.96 SEM) from 3 independent experiments. Different markers represent different experiments. Wild-type (WT): N=203 cells; mVenus-TERT (A67-A68): N=206 cells. Mann-Whitney U-test was performed.

### Simultaneous live-cell tracking of cell death and TERT localization

In addition to the evaluations of telomerase activity and localization, we compared the survival of cells harbouring wild-type TERT and mVenus-TERT to assess the possibility that mVenus compromises cell survival. We performed the following three simultaneous fluorescence measurements: dead cells visualized by SYTOX Blue staining, mitochondria visualized by MitoTracker Deep Red FM, and TERT visualized by mVenus (Fig. 4A). The time until apoptosis was calculated as the time when the fluorescence intensity of SYTOX Blue reached a specified threshold (see Methods Microscopy for live-cell imaging and tracking). Also, we calculated MCC of TERT with mitochondria from the fluorescence intensity at each time point (Fig. 4B), finding cells expressing wild-type TERT or mVenus-TERT had similar survival percentages after oxidative stress (Fig. 4C). This result showed that, by optimizing the insertion location, it is possible to produce a fluorescently labeled TERT that retains not only its normal localization and activity, but also normal cell-death properties upon oxidative stress. Moreover, live-cell imaging without oxidative stress did not show any cytotoxicity (Supplementary Fig. S2). Therefore, we concluded that cell death after oxidative stress was caused mainly by stress and not by mVenus.

**Figure 4.**
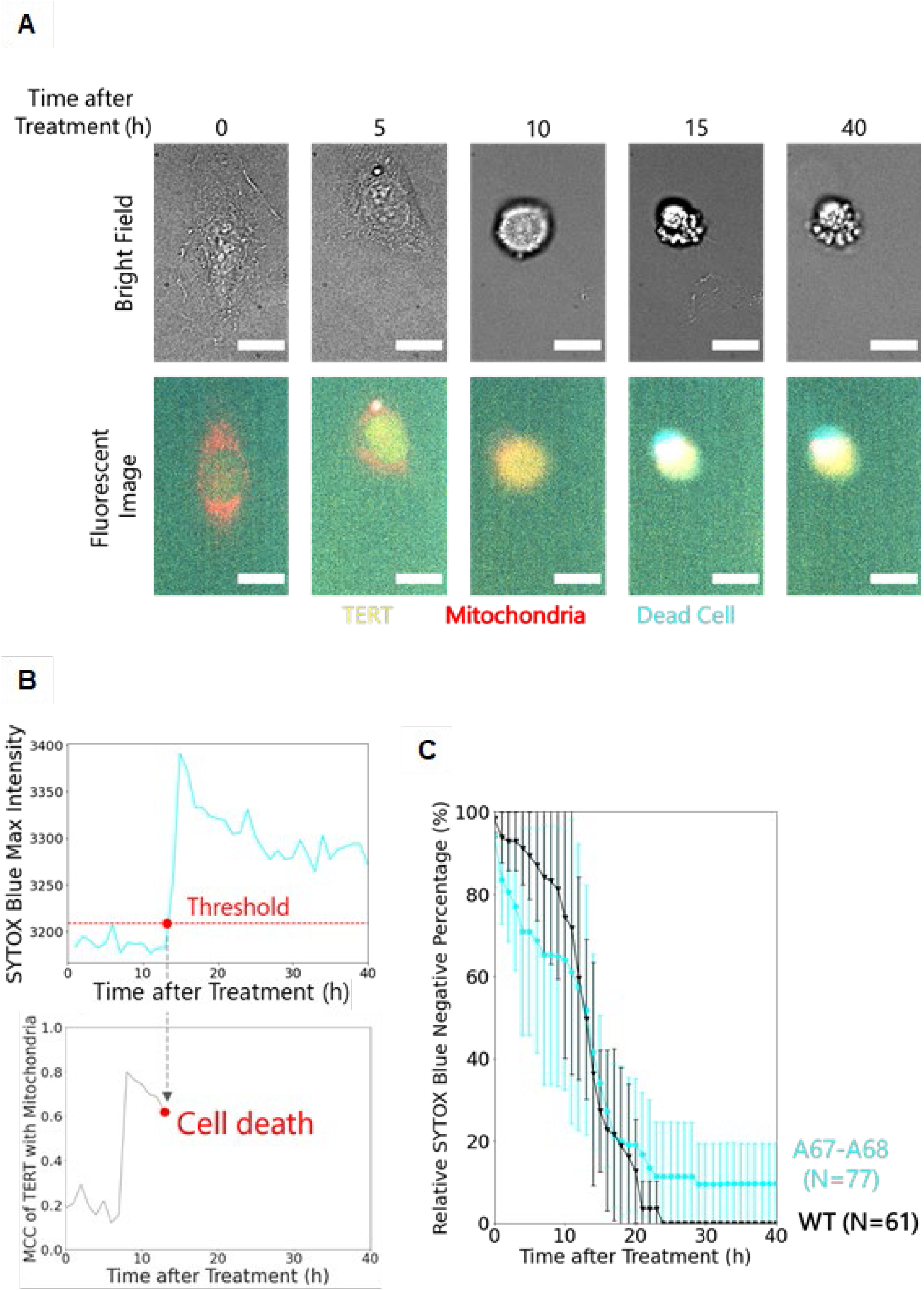
Simultaneous live-cell tracking of cell death and TERT localization. A. Representative live-cell images of HeLa cells expressing mVenus-TERT constructs. Yellow, mVenus fluorescence; red, MitoTracker Deep Red FM fluorescence; cyan, SYTOX Blue fluorescence. Cells were treated with 267 μM calcium percarbonate for 3 hours before the imaging. Scale bars, 20 μm. B. Representative time-course traces of SYTOX Blue fluorescence intensity and MCC of TERT with mitochondria of the cell in A. Cells were determined as positive after the SYTOX Blue fluorescence intensity exceeded the threshold. C. Percentages of SYTOX Blue-negative cells after oxidative stress among cells expressing wild-type TERT or mVenus-TERT. All percentages were corrected by control experiments without oxidative stress (Supplementary Fig. 2). Error bars show 95% C.I. (1.96 SEM) from 3 independent experiments. Wild-type (WT): N=61 cells; mVenus-TERT (A67-A68): N=77 cells.

### Live-cell tracking revealed opposing effects of mitochondrial TERT in apoptosis

From the live-cell tracking, we obtained the time-course plot of the MCC of TERT with mitochondria after oxidative stress in cells expressing mVenus-TERT. This plot demonstrated that cells with high MCC did not survive and all surviving cells showed low MCC (Fig. 5A). In contrast, cells cultured without oxidative stress did not show the cell population with high MCC (Supplementary Fig. S3). Apoptotic cells show unique morphological features, such as shrunk cellular bodies, meaning high MCC can result from the false detection of nuclear TERT as mitochondrial TERT. Live-cell tracking for the nuclear stain Hoechst demonstrated that cell death increased the MCC of Hoechst with mitochondria, which is consistent with the MCC of TERT with mitochondria (Supplementary Fig. S4). However, unlike MCC of TERT, MCC of Hoechst did not show an additional peak at high values, verifying that the high MCC of TERT with mitochondria is not an experimental artifact and that TERT actually accumulated in mitochondria upon oxidative stress in a subset of cells. Fitting the time-course plots of MCC showed that TERT barely changed its localization after oxidative stress (Supplementary Fig. S5E). Additionally, we found a positive correlation between MCC at the start of the imaging and time until apoptosis in cells expressing mVenus-TERT (Fig. 5B). This observation indicates that cells expressing wild-type TERT with high MCC tend to take longer to undergo apoptosis. From these results, we established a model for the role of mitochondrial TERT in apoptosis (Fig. 5C).

**Figure 5.**
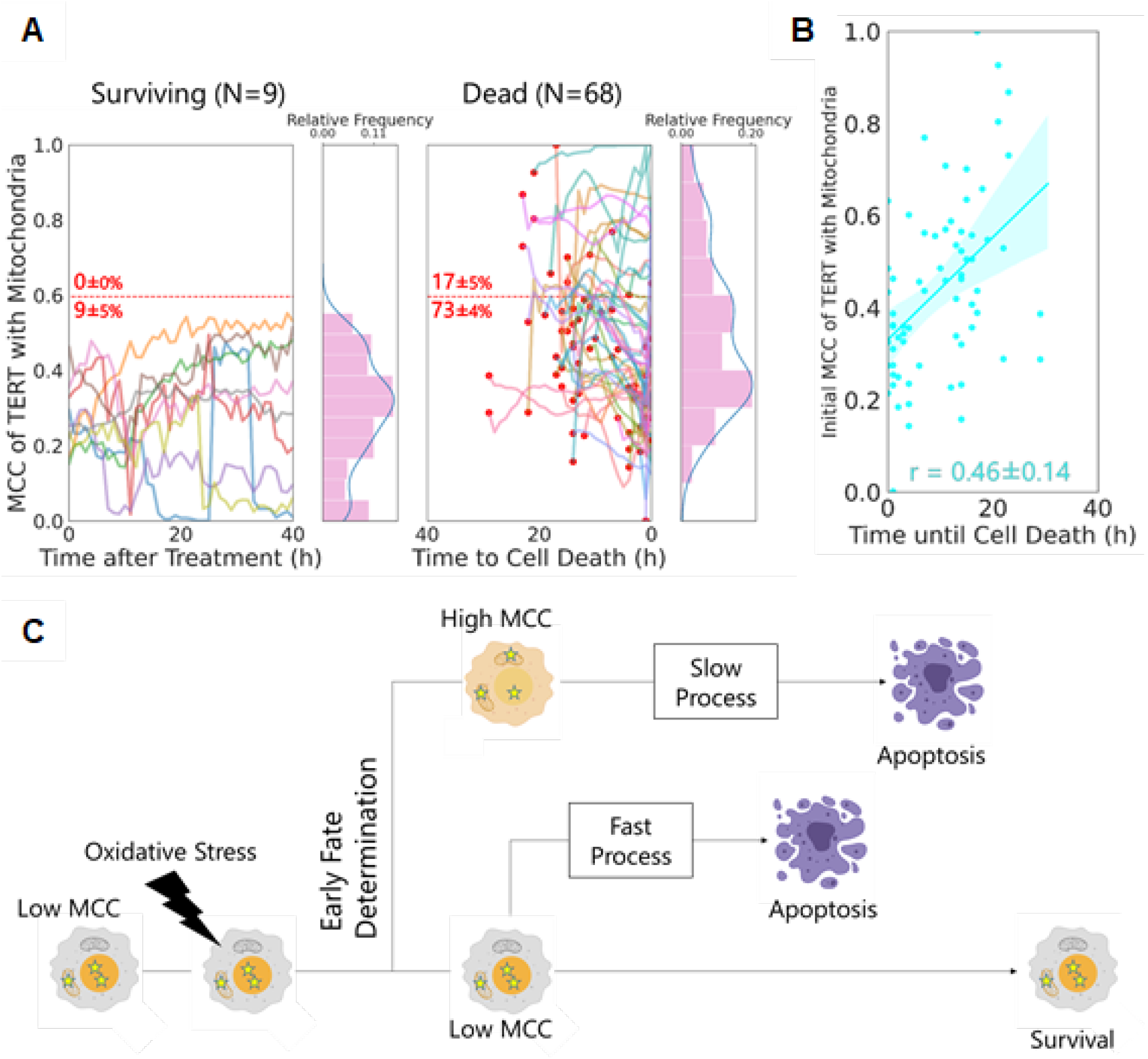
Live-cell tracking revealed the effects of mitochondrial TERT in apoptosis. A. Time-course plot of MCC of TERT with mitochondria in each cell. A histogram and kernel density estimation (KDE) plot of all MCC are shown. Plotted values are the mean values per 5 frames. For dead cells, 0 in the x-axis represents the moment the cells died. Red dots show MCC at the beginning of the observation. Red numbers show the percentage (mean ± SEM from 3 independent experiments) of cells whose MCC was above or below the threshold represented by the red dashed line. Surviving cells (Surviving): N=9 cells; Dead cells (Dead): N=68 cells. B. A scatter plot and regression line between the initial MCC of each dead cell and duration until cell death. Translucent bands around the regression line represent 95% C.I. (1.96 SEM). r, Pearson’s correlation coefficient (PCC), shows the mean ± SEM from 3 independent experiments. N=68 cells. C. A new model for the roles of mitochondrial TERT in apoptosis. This figure was created with BioRender.com.

### TERT mutants R3E/R6E and Y707F inhibited the death of cells with low MCC and delayed apoptosis independent of TERT localization

We also evaluated the effects of the TERT mutations using our live-cell tracking system. Similar to the live-cell imaging of wild-type TERT, TERT mutations R3E/R6E and Y707F increased cell survival even if bound to mVenus (Supplementary Fig. S5A). Also, the time course of MCC of TERT with mitochondria after oxidative stress in these cells showed that almost all cells with high MCC died by oxidative stress, consistent with mVenus-TERT (Supplementary Fig S5B). However, unlike mVenus-TERT, there was no correlation between the initial MCC and time until apoptosis in the cells expressing mVenus-TERT mutants (Fig. 5B and Supplementary Fig. S5C). Next, to visualize the difference between the cells with high or low initial MCC, we set a threshold at approximately 0.60 (red dashed lines in Fig. 5A and Supplementary Fig. S5B), as only a small subset of surviving cells showed higher initial MCC than the threshold (0% for mVenus-TERT, 0% for mVenus-TERT R3E/R6E and 2% for mVenus-TERT Y707F). This division indicated that mVenus-TERT mutants caused a lower percentage of dead cells having low MCC (Supplementary Fig. S5B). The time until the apoptosis of cells with low MCC of mVenus-TERT mutants was the same as those with high MCC mVenus-TERT (Supplementary Fig. S5D), explaining no correlation between the initial MCC and time until apoptosis in the cells expressing mVenus-TERT mutants. In addition, the time until the apoptosis of cells expressing mVenus-TERT was equivalent to that of cells with high MCC of mVenus-TERT and longer than that of cells with low MCC of mVenus-TERT. The time difference between mVenus-TERT and mVenus-TERT mutants in cells with low MCC indicates that the mutants delayed cell death induced by oxidative stress. Taken together, the mutations showed inhibited and delayed effects on stress-induced cell death in cells showing low MCC.

## DISCUSSION

To reveal the role of mitochondrial TERT in apoptosis, it is essential to observe the relationship between the TERT distribution in each cell and cell fate. Here, we achieved this by combining live-cell imaging-based dead-cell detection with the observation of a new mVenus-TERT reporter, which retains TERT properties. Our simultaneous tracking of the cell-death process and TERT distribution of individual cells demonstrated a strong correlation between a high mitochondrial accumulation of TERT and cell death, but the high accumulation was also positively correlated with a longer time until cell death. These results suggest that TERT localization in mitochondria plays distinct roles at different stages of apoptosis.

Based on these results, we propose a new model that integrates the seemingly contradictory data from previous studies for the role of TERT in apoptosis. We speculate that there are two stages after oxidative stress. In the first stage, which is immediately after oxidative stress, mitochondrial TERT promotes apoptosis. Live-cell tracking of wild-type TERT localization revealed that after oxidative stress, all cells with high MCC of TERT with mitochondria died, while all surviving cells showed lower MCC. This observation of the mitochondrial localization of TERT after oxidative stress is consistent with previous reports [4, 5]. In the present study, the cells experienced 3 hours of oxidative stress before tracking, and cells with high MCC appeared in the first frame of the tracking. Therefore, cells that showed an accumulation of mitochondrial TERT had their fate determined within 3 hours of the oxidative stress treatment. Intrinsic apoptosis is triggered by mitochondrial outer membrane permeabilization (MOMP), and thus, mitochondrial accumulation of TERT can result from passive flux of TERT into mitochondria that undergo MOMP, leading to the outcome that all cells with high mitochondrial accumulation of TERT result in apoptosis. However, mitochondrial translocation of TERT has been reported to depend on mitochondrial membrane potential (MMP) and TERT did not accumulate to mitochondria when cells were treated with valinomycin, which compromise MMP [18]. Although valinomycin treatment may be not necessarily analogous to MOMP, the onset of MOMP is often associated with a loss of MMP [23]. Thus, it is likely that after oxidative stress, TERT actively accumulates to mitochondria to predetermine apoptosis. Because all the cells with high MCC died after oxidative stress in this study and other reports found mitochondrial TERT induces apoptosis [4, 6], we propose a model in which this accumulation predetermines apoptosis.

However, in the second stage, mitochondrial TERT delays the apoptotic process. In cells expressing wild-type TERT, the initial MCC of TERT with mitochondria positively correlated with the time until apoptosis. This correlation indicates that more TERT in mitochondria delays apoptosis. This delay can explain why previous studies found that mitochondrial TERT suppresses apoptosis [5, 7]. Several reports suggest that TERT inhibits mitochondrial pathway of apoptosis through the interaction with anti-apoptotic protein Bcl-2, such as inhibiting the conformational activation of pro-apoptotic Bcl-2 family protein BAX [24, 25]. TERT has also been reported to have a conserved motif among Bcl-2 family proteins and to interact with the anti-apoptotic Bcl-2 family proteins Bcl-xL and Mcl-1 via the motif [26]. The live-cell tracking method that we developed in this study will reveal the real-time interaction between TERT and Bcl-2 family proteins during apoptosis.

Both TERT mutations R3E/R6E and Y707F are thought to suppress apoptosis by inhibiting the mitochondrial translocation of TERT. We evaluated this possibility by live-cell tracking. Similar to wild-type TERT, almost all cells with high MCC of TERT with mitochondria did not survive, demonstrating that these mutations did not completely inhibit the mitochondrial translocation of TERT. Further, the mutations did not inhibit the mitochondrial localization of TERT without oxidative stress. These results are in contrast to previous reports, which evaluated the effect of these mutations from representative immunofluorescence and Western blotting data without quantification [4, 6, 27]. We, on the other hand, quantified the TERT localization in each cell by a colocalization index, MCC. Furthermore, we evaluated the localization of TERT mutant R3E/R6E in HeLa cells, which highly express TERT endogenously, whereas previous studies evaluated the localization of this TERT mutant in normal human fibroblasts and MRC-5 cell lines, both of which do not express TERT endogenously. Since TERT oligomerization has been reported [28], it is possible that in our study, endogenous wild-type TERT oligomerized with exogenous TERT carrying R3E/R6E for transport to mitochondria. To test this hypothesis, knockout experiments of the TERT locus from HeLa cells would be helpful. In any case, our data suggest that TERT mutations R3E/R6E and Y7070F suppress apoptosis in cells without TERT accumulating in mitochondria. Therefore, we assume that these mutations suppress apoptosis independently of mitochondrial TERT accumulation.

Compared with mVenus-TERT, mVenus-TERT mutations R3E/R6E and Y707F suppressed and delayed apoptosis in the cells with low MCC, suggesting that these mutations alter the cell death capacity of nuclear TERT under oxidative stress. Regardless of its mitochondrial accumulation, TERT has been reported to act as a suppressor of apoptosis [29, 30]. The negative effects of the TERT mutations on stress-induced cell death are likely due to such a role of TERT in pathways other than mitochondrial. For instance, besides its canonical function as a telomerase, TERT has been reported to serve as an RNA-dependent RNA polymerase (RdRP) and to regulate siRNAs and miRNAs [31, 32]. In addition, TERT has also been reported to act as a transcription regulator through the pathway such as Myc [33, 34], Wnt/β-catenin [33, 35], and NF-κB [36]. Especially, NF-κB pathway regulate apoptosis-related proteins such as Bcl-2. R3E/R6E and Y707F might change the TERT capacity as a transcription regulator and the expression pattern of apoptosis-related proteins, resulting in the blunted effect of Z-VAD-fmk to these mutants. The investigation of the gene expression in cells expressing these mutants and the transcription pathways of the cells will reveal the effect of these TERT mutations on the apoptotic pathway.

The live-cell tracking method that we developed in this study is a potential gold standard for investigating the effect of proteins on cell death. Applying this method to other cell lines besides HeLa cells will elucidate whether our finding is universal. In the current system, accurate subcellular imaging is limited to adherent cells, but combining live-cell tracking tiny microwells [37] or flow cytometry [38] will broaden the method to floating cells. Also, live-cell tracking can be used to investigate other biological phenomena that are related to mitochondrial TERT, such as DNA protection [4, 20], ROS production [4, 7, 19], autophagy [27], mitophagy [39], and senescence [6, 40, 41]. In addition, apoptosis has several different pathways even within the same cell line [8, 9]. Thus, to unravel the molecular basis of mitochondrial TERT in apoptosis, it is crucial to obtain the state of each individual cell before interactome and transcriptome analysis. Our method can provide such cellular information and thus can contribute to future studies on the molecular basis of mitochondrial TERT, which will accelerate our understanding of how TERT participates in the metabolism of cancer cells beyond its function as a telomerase.

## MATERIALS AND METHODS

### Plasmids

The gene encoding TERT was amplified from the plasmid pCDH-3xFLAG-TERT, which was a gift from Steven Artandi (Addgene plasmid #51631; http://n2t.net/addgene:51631). The mVenus sequence was isolated from pCS2-mVenus plasmid, which was purchased from the RIKEN BioResorce Research Center (RDB15116, Ibaraki, Japan). pSBbi-Pur was a gift from Eric Kowarz (Addgene plasmid #60523; http://n2t.net/addgene:60523). The gene encoding TERT and mVenus was inserted into pSBbi vector using the In-Fusion cloning kit (Takara Bio, Shiga, Japan). The R3E/R6E mutation was introduced using the KOD mutagenesis kit (Toyobo, Osaka, Japan). The Y707F mutation was introduced by PCR. All plasmids were transformed into DH5α chemical competent cells and purified using the FastGene miniprep kit and endotoxin-free miniprep kit (NIPPON Genetics, Tokyo, Japan). The primer sets used are presented in Supplementary Table S1.

### Cell culture and generation of stable cell lines

The HeLa cell line was purchased from the RIKEN BioResource Research Center (RCB0007). Cells in 6-well plates were transfected with 1.9 µg of pSBbi-TERT-Pur plasmid, 0.1 µg of pCMV-SB100 [10] and 3.75 µg of polyethylenimine in 250 µL of Opti-MEM. After transfection, cells were selected in DMEM supplemented with 10% FBS, 1 × PenStrep, and 1 µg/mL Puromycin for one week. All cells were cultured in DMEM supplemented with 10% FBS and 1 × PenStrep in a 37°C incubator at 5% CO_2_.

### Immunofluorescence

Two days before immunofluorescence, the cells were passaged in a black wall poly-L-lysine-coated multi-well glass bottom dish (Matsunami, Osaka, Japan) at a density of 4000 cells per well. The cells were incubated with DMEM supplemented with 10% FBS, 1 × PenStrep, and 100 nM MitoTracker Deep Red FM (Thermo Fisher Scientific, Waltham, MA USA) for 30 minutes at 37°C and 5% CO_2_. All procedures after incubation with MitoTracker Deep Red FM were performed under room temperature in the dark. The washing step referred to exchange the medium for PBS (Nacalai, Kyoto, Japan) and to incubate the cells for 5 minutes.

Fixation was performed using 4% paraformaldehyde in PBS for 10 min followed by washing. Permeabilization was performed with 0.5% Triton-X-100 in PBS for 10 min followed by 3 washes. Blocking was performed with 3% BSA in PBS-T for 1 hour followed by washing.

For immunofluorescence with the TERT mutations, the cells were incubated with 20 µM Cellstain Hoechst 33342 (Wako, Osaka, Japan) in PBS for 10 minutes and washed. The cells were then incubated with anti-TERT antibody (Rockland, Limerick, PA USA, 600-401-252S, 1:500 dilution) in PBS-T for 1 hour and washed 3 times. Finally, the cells were incubated with anti-rabbit IgG (H+L), F(ab’)2 fragment conjugated with Alexa Fluor 488 (CST, Danvers, MA USA) in PBS-T for 1 hour and washed 3 times.

For the immunofluorescence of cells expressing mVenus-TERT, after blocking, the cells were washed 3 times and then incubated with anti-TERT antibody conjugated with CF405M (1:100 dilution) in PBS-T for 1 hour and washed 3 times. Dye conjugation was performed using the Mix-n-Stain CF405M Antibody Labeling Kit (Biotium, Fremont, CA USA). Anti-TERT antibody conjugated with CF405M was diluted at 1:5 in storage buffer.

The imaging medium for immunofluorescence was PBS supplemented with 1% ProLong Live Antifade Reagent (Thermo Fisher Scientific).

Immunofluorescence imaging of the cells was performed using a SpinSR10 (Olympus, Tokyo, Japan) with an oil-immersion objective (PlanApoN 60×/1.40 Oil, Olympus). Fluorophores were excited at 405 nm (for Hoechst or CF405M), 488 nm (for Alexa Fluor 488), 512 nm (for mVenus), and 640 nm (for MitoTracker Deep Red FM).

### Dead-cell detection with live-cell imaging

Cells were cultivated for a week before the assay and passaged 5 times. A day before the assay, the cells were passaged into black wall poly-L-lysine coated 96-well plates at a density of 1000 cells per well. All empty wells and space between wells were filled with sterilized water.

Cells were treated with DMEM supplemented with 10% FBS, 1 × PenStrep, and 267 μM sodium carbonate hydrogen peroxide (equivalent to 267 μM sodium carbonate and 400 μM hydrogen peroxide) for 3 hours. As negative controls, cells were treated with DMEM supplemented with 10% FBS, 1 × PenStrep, and 267 μM sodium carbonate. During imaging, all cells were cultured in DMEM supplemented with 10% FBS, 1 × PenStrep, 250 nM SYTOX Orange (Thermo Fisher Scientific) and 2.5 μM YO-PRO-1 (Thermo Fisher Scientific) at 37°C and 5% CO_2_.

For live-cell imaging with Z-VAD-fmk, the cells were treated with DMEM supplemented with 10% FBS, 1 × PenStrep, 50 μM Z-VAD-fmk, and 267 μM sodium carbonate hydrogen peroxide for 3 hours. Then, the cells were cultured in DMEM supplemented with 10% FBS, 1 × PenStrep, 50 μM Z-VAD-fmk, 250 nM SYTOX Orange (Thermo Fisher Scientific) and 2.5 μM YO-PRO-1 (Thermo Fisher Scientific) at 37°C and 5% CO_2_.

### Telomerase activity assay

5.0 × 10^6^ cells were harvested and frozen at -80°C. Thawed cells were treated with and assayed following the instructions of the Telomerase Activity Quantification qPCR Assay Kit (ScienCell, Carlsbad, CA USA). qPCR was performed using OneStepPlus (Thermo Fisher Scientific).

### Dead-cell detection and TERT visualization by live-cell tracking

Cells were cultivated a week before the assay and passaged 5 times. A day before the assay, the cells were passaged into black wall poly-L-lysine coated 96-well plates at a density of 1000 cells per well. All empty wells and space between wells were filled with sterilized water. Cells were incubated with DMEM supplemented with 10% FBS, 1 × PenStrep, and 100 nM MitoTracker Deep Red FM for 30 minutes at 37°C and 5% CO_2_. For the oxidative stress, the cells were treated with DMEM supplemented with 10% FBS, 1 × PenStrep, and 267 µM sodium carbonate hydrogen peroxide (equivalent to 267 μM sodium carbonate and 400 μM hydrogen peroxide) for 3 hours. As negative controls, cells were treated with DMEM supplemented with 10% FBS, 1 × PenStrep, and 267 μM sodium carbonate. During imaging, all cells were cultured in DMEM supplemented with 10% FBS, 1 × PenStrep and 250 nM SYTOX Blue (Thermo Fisher Scientific) at 37°C and 5% CO_2_.

For control experiments with Hoechst, cells were incubated with 10% FBS, 1 × PenStrep, and 100 nM MitoTracker Deep Red FM (Thermo Fisher Scientific) for 30 minutes at 37°C and 5% CO_2_, and then were incubated with 10% FBS, 1 × PenStrep, and 20 µM Cellstain Hoechst 33342 (Wako) for 15 minutes at 37°C and 5% CO_2_.

### Microscopy for live-cell imaging and tracking

Imaging was performed using an inverted microscope (Ti, Nikon, Tokyo, Japan) with an objective (Plan Apo Lambda 40×/0.95, Nikon), LED illumination system (X-Cite XLED1, Lumen Dynamics, Ontario, Canada) and filter sets [CFP-2432C (for SYTOX-Blue or Cellstain Hoechst 33342; Semrock), GFP-3035D (for YO-PRO-1; Semrock), LF514-B (for mVenus; Semrock), TRITC-A-Basic (for SYTOX-Orange; Semrock), and Cy5-4040C (for MitoTracker Deep Red FM; Semrock)]. Bright-field and fluorescence images were captured using an electron multiplying CCD camera (ImagEM X2-1K EM-CCD, Hamamatsu Photonics, Shizuoka, Japan). The sample temperature was kept at 37°C by feedback from a heat sensor in a water-filled well and was monitored by NECO (TOKAI HIT, Shizuoka, Japan).

For the live-cell imaging, dead cells were defined as cells that were stained with SYTOX Orange or YO-PRO-1; the threshold was set to the mean intensity of negative control cells plus 10 times the standard deviation using NIS-Elements software (Nikon).

For live-cell tracking, a 1.5× magnification lens was used, and dead cells were defined as cells that were stained with SYTOX Blue; the threshold was set to the mean intensity of negative control cells plus 5 times the standard deviation using NIS-Elements software.

For control experiments with Hoechst, dead cells were defined as cells that were stained with SYTOX Orange; the threshold was set to the mean intensity of negative control cells plus 10 times the standard deviation using NIS-Elements software.

### Colocalization analysis

Each cell in each frame in a bright field image was used as a reference to make regions of interest manually. Otsu’s method [35] was employed to find pixels above a certain fluorescence threshold, and Manders’ colocalization coefficients (MCCs) were calculated in the pixels using the GDSC colocalization plugins for Fiji/ImageJ.

MCC of TERT with mitochondria was calculated from TERT intensity *T* and mitochondria intensity *M* as:

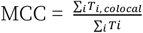

where *T*_*i, colocai*_ = *T*_*i*_ if *M*_*i*_ > 0 and *T*_*i, colocal*_ = 0 if *M*_*i*_ = 0.

Kernel density estimation (KDE) [43] was performed on MCC histograms in Fig. 5A and Supplementary Fig. S3, S4, and S5B. KDE estimates the true probability density function from the dataset, which corresponds to MCC distributions in this work.

### Statistical analysis

Statistical tests were performed using R software. The Steel-Dwass test was performed in Fig. 1B and 3A and Supplementary Fig. S5E using the NSM3 library with the Monte Carlo method. The Mann-Whitney U-test was performed in Fig. 3D.

## ACKNOWLEDGEMENTS

We thank Dr. Yoshitaka Shirasaki for critical advice, Uemura lab members for assistance with the experiments, and Dr. Keigo Ikezaki and Dr. Sawako Enoki for support with the confocal microscopy.

## AVAILABILITY OF DATA AND MATERIALS

The authors declare that the data are available within the paper and its Supplementary information files.

## AUTHOR CONTRIBUTIONS

Conceptualization, H.E. and T.S.; Methodology, H.E. and T.S.; Investigation, H.E.; Analysis, H.E., Writing – Original draft, H.E., T.S. and S.U; Draft editing, H.E., T.S., R.I., S.U.

## ETHICS

Our study did not require ethical approval.

## Supplemental Information

**Supplementary Figure S1.**
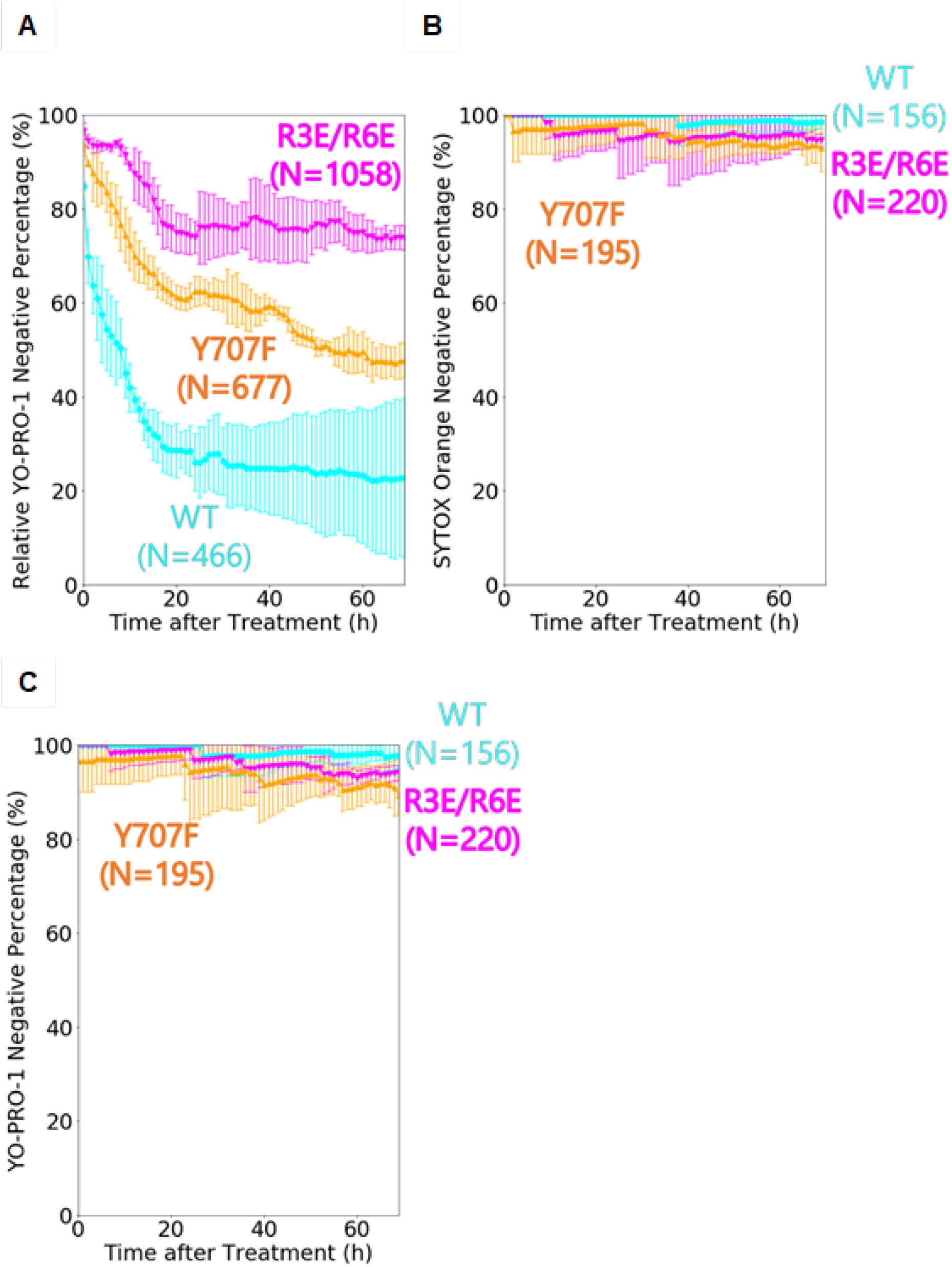
Survival curves of YO-PRO-1 after oxidative stress and of SYTOX Orange and YO-PRO1 without oxidative stress. A. The percentage of cells determined as YO-PRO-1-negative after oxidative stress. All percentages were corrected with the data in C. Error bars show 95% C.I. (1.96 SEM) from 3 independent experiments. Wild-type (WT): N=466 cells; R3E/R6E: N=1058 cells; Y707F: N=677 cells. B. The percentage of cells determined as YO-PRO-1-negative without oxidative stress. Error bars show 95% C.I. (1.96 SEM) from 3 independent experiments. Wild-type (WT): N=156 cells; R3E/R6E: N=220 cells; Y707F: N=195 cells. C. The percentage of cells determined as YO-PRO-1-negative without oxidative stress. Error bars show 95% C.I. (1.96 SEM) from 3 independent experiments. Wild-type (WT): N=156 cells; R3E/R6E: N=220 cells; Y707F: N=195 cells.

**Supplementary Figure S2.**
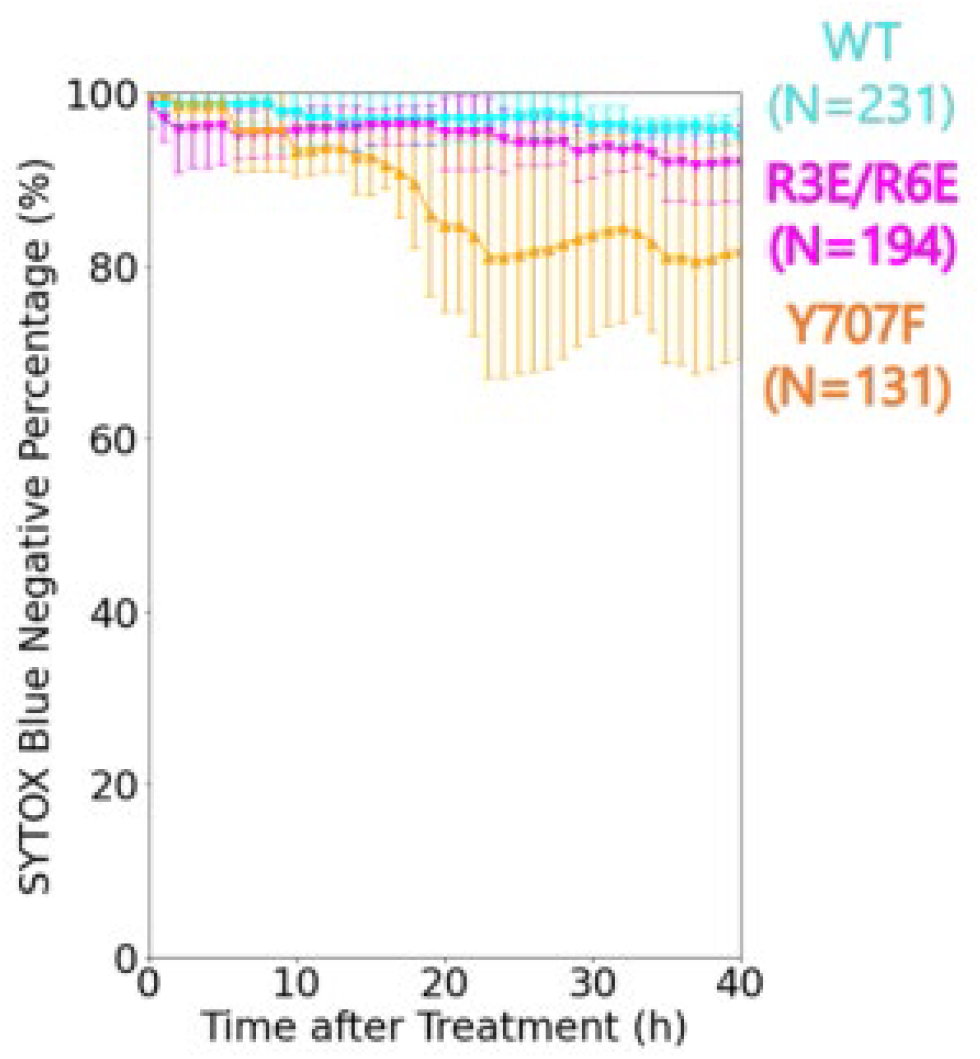
Imaging did not show cytotoxicity without oxidative stress. The percentage of cells determined as SYTOX Blue-negative without oxidative stress. Error bars show 95% C.I. (1.96 SEM) from 3 independent experiments. mVenus-TERT (WT): N=231 cells; mVenus-TERT R3E/R6E (R3E/R6E): N=194 cells; mVenus-TERT Y707F (Y707F): N=131 cells.

**Supplementary Figure S3.**
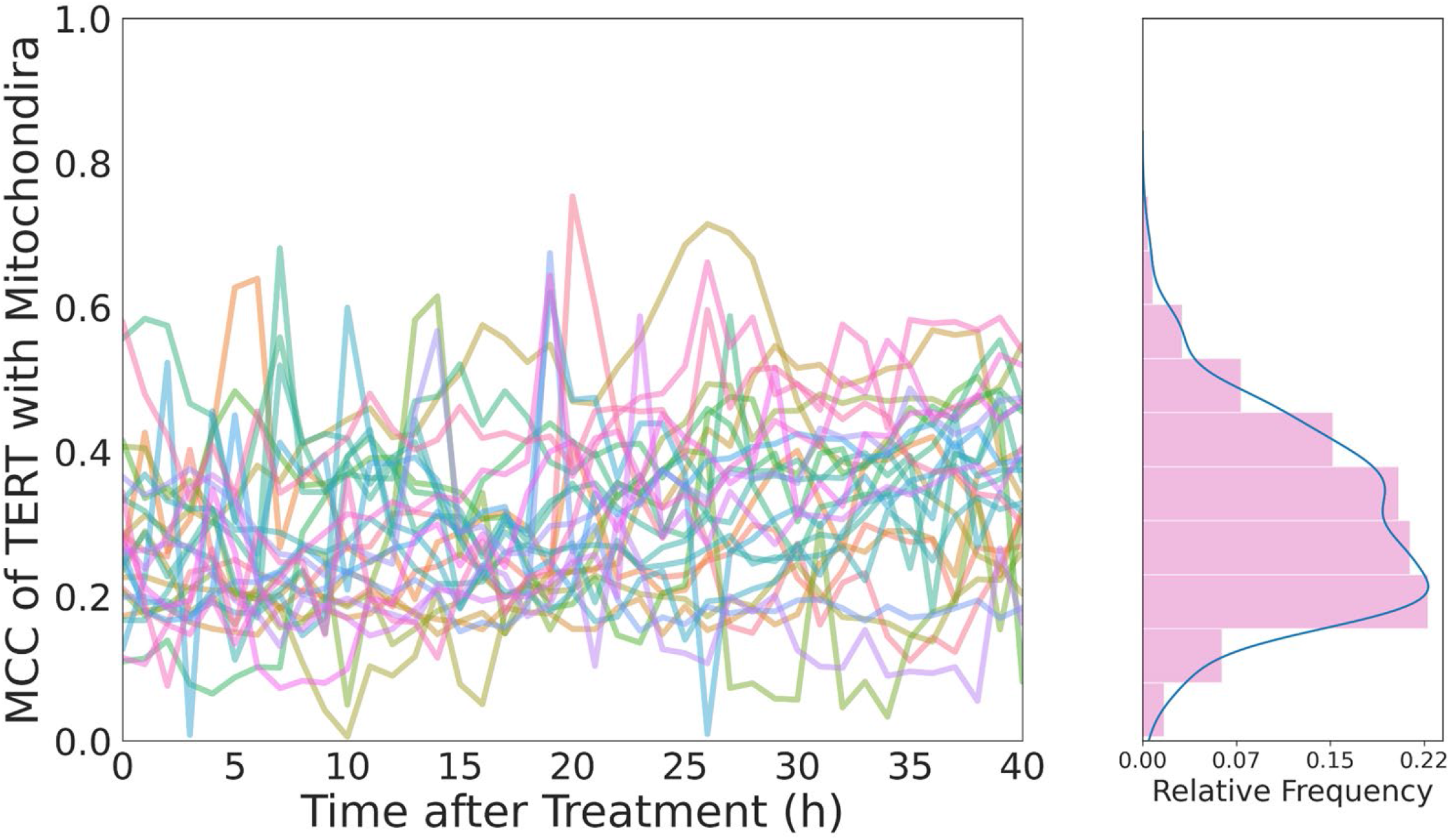
Cells without oxidative stress did not show high MCC of TERT with mitochondria. Histogram and KDE plot of MCC of TERT with mitochondria of cells incubated without oxidative stress. Plotted values are the mean of values per 5 frames. N= 27 cells.

**Supplementary Figure S4.**
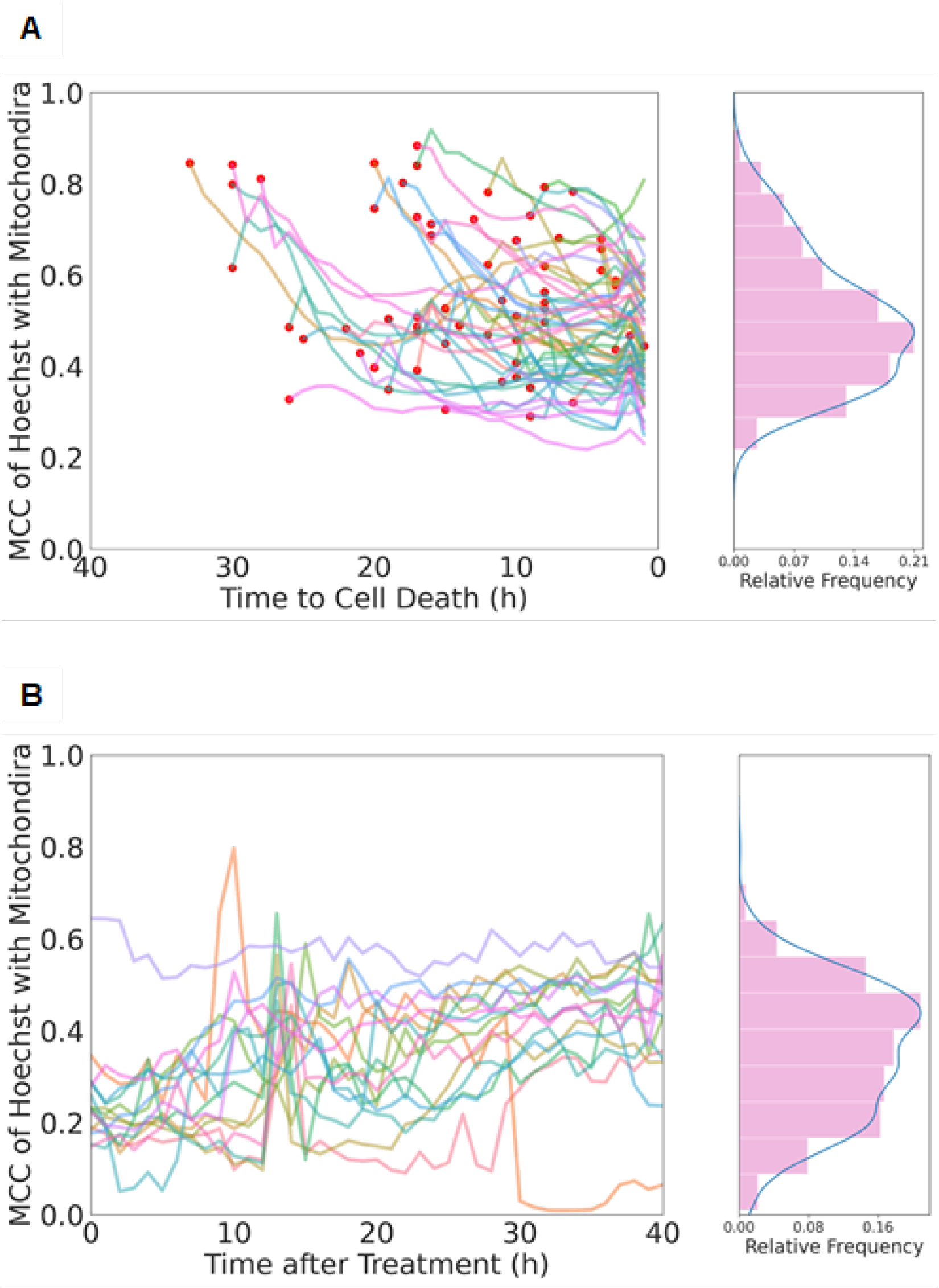
Dead cells did not show high MCC of Hoechst with mitochondria after oxidative stress. A. Histogram and KDE plot of MCC of Hoechst with mitochondria of cells after oxidative stress treatment. Plotted values are the mean of values per 5 frames. N=61 cells. Red dots show the MCC of Hoechst with mitochondria at the beginning of the observation. B. Histogram and KDE plot of MCC of Hoechst with mitochondria of cells without oxidative stress. Plotted values are the mean of values per 5 frames. N=16 cells.

**Supplementary Figure S5.**
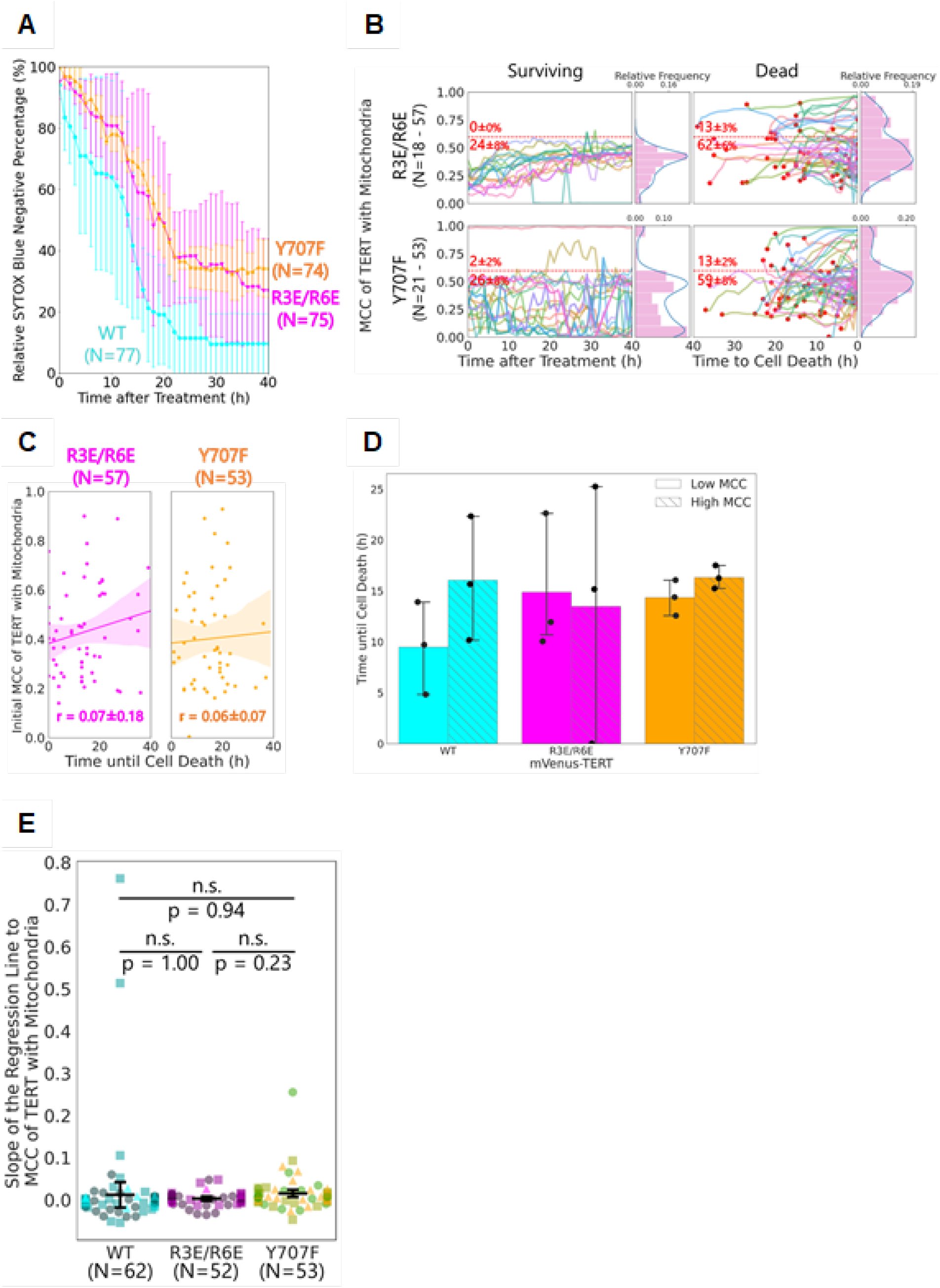
TERT mutants R3E/R6E and Y707F decreased the apoptosis of cells with low MCC and lost the positive correlation between the initial MCC and time until cell apoptosis. A. The percentage of cells determined as SYTOX Blue-negative after oxidative stress. All percentages were corrected by control cells (without oxidative stress; see Supplementary Fig. 3B). Error bars show 95% C.I. (1.96 SEM) from 3 independent experiments. mVenus-TERT (WT): N=77 cells; mVenus-TERT R3E/R6E (R3E/R6E): N=75 cells; mVenus-TERT Y707F (Y707F): N=74 cells. B. Histogram and KDE plot of MCC of TERT with mitochondria in each cell. Plotted values are the mean of values per 5 frames. For dead cells, the beginning of cell death was set to 0 in the x-axis. Red dots show MCC of TERT with mitochondria at the beginning of the observation. Red numbers show the percentage (mean ± SEM from 3 independent experiments) of cells whose initial MCC of TERT with mitochondria was above or below the threshold represented by the red dashed line. For mVenus-TERT R3E/R6E (R3E/R6E), surviving cells (Surviving): N=18 cells; Dead cells (Dead): N=57 cells. For mVenus-TERT Y707F (Y707F), surviving cells (Surviving): N=21 cells; Dead cells (Dead): N=53 cells. C. Scatter plot and regression line between the initial MCC of TERT with mitochondria of each dead cell and time until cell death. Translucent bands around the regression line represent 95% C.I. (1.96 SEM). r, Pearson’s correlation coefficient (PCC), shows the mean ± SEM from 3 independent experiments. mVenus-TERT R3E/R6E (R3E/R6E): N=57 cells; mVenus-TERT Y707F (Y707F): N=53 cells. D. Time until death of cells with high or low initial MCC of TERT with mitochondria. Graphs show data from each cell and the mean ± 95% C.I. (1.96 SEM) from 3 independent experiments. E. Slope of the regression line of MCC of TERT with mitochondria. Dots show the slope of each dead cell, and bars show the mean ± 95% C.I. (1.96 SEM) from 3 independent experiments. Different markers represent different experiments. mVenus-TERT (WT): N=62 cells; mVenus-TERT R3E/R6E (R3E/R6E): N=52 cells; mVenus-TERT Y707F (Y707F): N=53 cells. Steel-Dwass test was performed.

**Supplementary Table S1.**
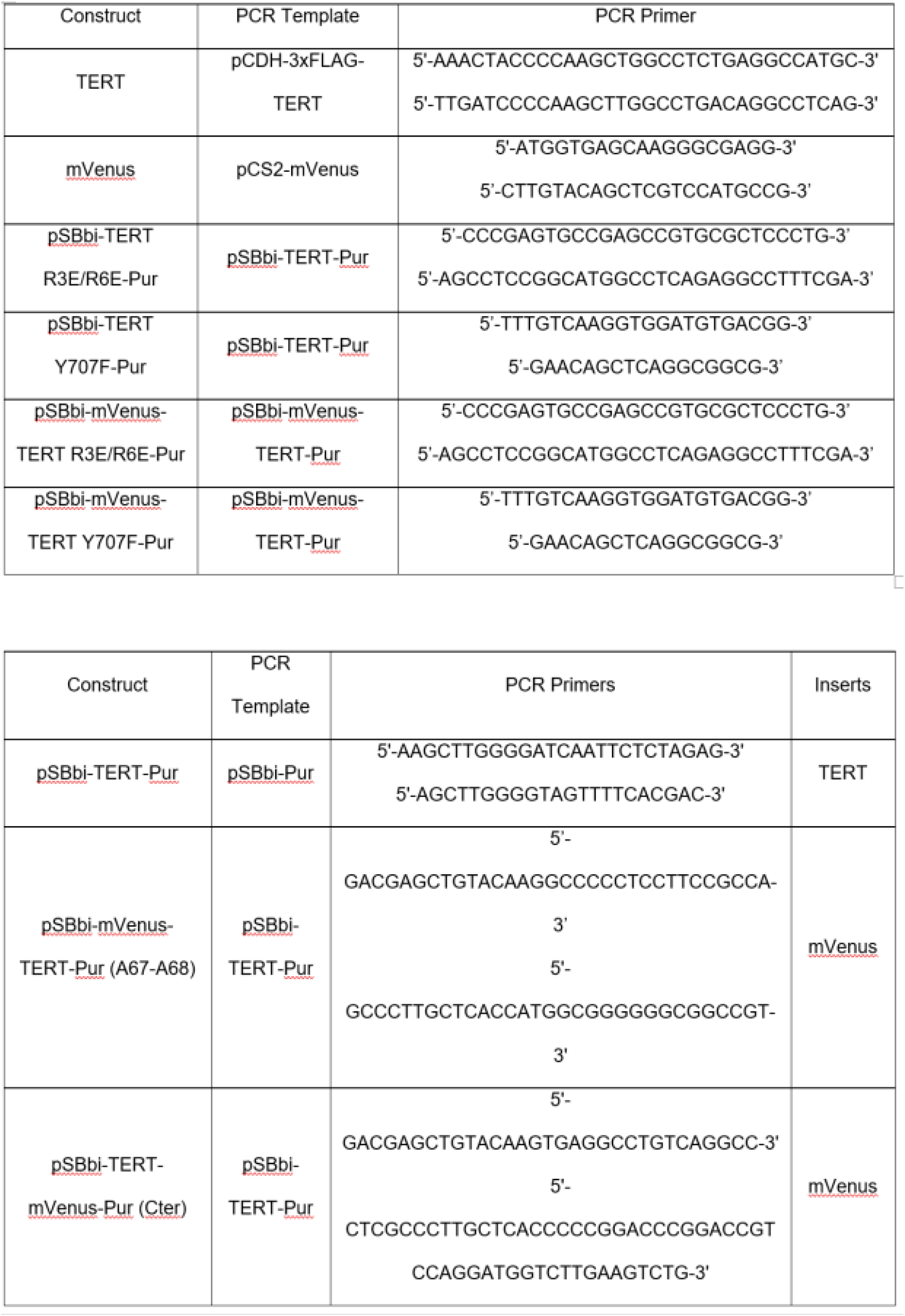
Key PCR primers used to generate the TERT constructs in this study.

